# One-dimensional sliding assists σ^70^-dependent promoter binding by Escherichia coli RNA polymerase

**DOI:** 10.1101/494534

**Authors:** Iddo Heller, Margherita Marchetti, Abhishek Mazumder, Anirban Chakraborty, Agata M. Malinowska, Wouter H. Roos, Richard H. Ebright, Erwin J.G. Peterman, Gijs J.L. Wuite

## Abstract

The search for a promoter on DNA by RNA polymerase (RNAP) is an obligatory first step in transcription. The role of facilitated diffusion during promoter search has been controversial. Here, we re-assessed facilitated diffusion in promoter search by imaging motions of single molecules of *Escherichia coli* RNAP σ^70^ holoenzyme on single DNA molecules suspended between optical traps in a manner that absolutely avoided interactions with surfaces. The assay enabled us to observe unambiguous one-dimensional sliding of RNAP σ^70^ holoenzyme for thousands of DNA base pairs during promoter search. Analysis of binding kinetics revealed short binding events on nonspecific DNA (0.4 s), intermediate binding events on A/T-rich DNA (1.6 s), and long binding events at or near promoters (>300 s). We estimate a *lower bound* for the “diffusion facilitation threshold” – the RNAP concentration at which three-dimensional search and one-dimensional sliding contribute equally to promoter binding – of 0.2 μM RNAP. The results suggest facilitated diffusion occurs in promoter search by RNAP, even at the relatively high, 0.2-0.6 μM, concentrations of RNAP in cells.

**Significance statement:** The flow of genetic information from DNA to RNA is of central importance to living systems, and it can only start after an RNA polymerase (RNAP) has found a promoter site. But how does this enzyme find promoter sites on DNA in the first place? In recent years, debate on this topic has favored a promoter search mechanism that is dominated by three-dimensional diffusion of RNAP, rather than by one-dimensional sliding of RNAP on DNA. Here, we designed an improved single-molecule assay that unambiguously revealed extensive one-dimensional sliding of RNAP on DNA. Our results imply that, at the RNAP concentrations in living cells, the promoter-search process is facilitated by one-dimensional sliding on DNA.

## Introduction

In living cells, the enzyme RNA polymerase (RNAP) is responsible for RNA synthesis. Bacterial RNAP holoenzyme comprises RNAP core enzyme, which catalyzes RNA polymerization, and the transcription initiation factor σ, which mediates recognition of specific DNA sites, termed “promoters,” preceding genes. The rate of promoter binding is an obvious target for regulation of gene expression. This rate is determined by the “promoter-search” mechanism, which, in principle, may include three-dimensional (3D) molecular motions through bulk solvent, one-dimensional (1D) molecular motions along DNA, or both (1–3).

The mechanism of promoter search mechanism by RNAP holoenzyme has been controversial (4–7). Early studies, conducted both at the ensemble level and at the single-molecule level, reported that RNAP holoenzyme can move long distances along DNA by 1D sliding (2, 4, 5, 8, 9), supporting the notion that facilitated diffusion contributes to promoter search. However, recent studies by Wang et al. and Friedman et al. (6, 10), conducted at the single-molecule level, observed little or no 1D sliding. The study of Wang et al. entailed imaging movements of *Escherichia coli* RNAP holoenzyme containing the primary σ factor, σ^70^, responsible for transcription initiation at most promoters, on surface-bound “DNA curtains” (6); the study of Friedman et al. entailed fluorescence-colocalization imaging (CoSMoS) of *E. coli* RNAP holoenzyme containing the structurally and mechanistically distinct alternative σ factor, σ^54^, which is responsible for transcription initiation at promoters of genes required for response to nitrogen starvation (10). The difference between the results of the early studies and the results of Friedman et al. (10) readily can be explained by the analysis of different *E. coli* RNAP holoenzyme derivatives: the early studies analyzed RNAP holoenzyme containing the primary σ factor, σ^70^, whereas Friedman et al. analyzed RNAP holoenzyme containing the alternative σ factor, σ^54^ (10). In contrast, the difference between the results of the early studies and the results of Wang et al. (6) is difficult to explain, but potentially could reflect artefactual restrictions on DNA sliding by RNAP holoenzyme arising from non-native interactions between DNA and surfaces, and/or non-native interactions between RNAP holoenzyme and surfaces, in the surface-bound DNA curtains analyzed in Wang et al (6).

In this work, we sought to elucidate definitively the mechanism of promoter search by *E. coli* RNAP σ^70^ holoenzyme. We employed a single-molecule approach that enabled real-time, direct investigation of binding kinetics, binding locations, and mobility of individual RNAP σ^70^ holoenzymes on DNA and that, crucially, completely avoided non-native interactions between DNA and surfaces and non-native interactions between RNAP holoenzyme and surfaces. We observe unambiguous 1D sliding of RNAP-σ^70^ holoenzyme on DNA – 1D sliding for thousands of DNA base pairs – during promoter search. Our results establish that facilitated diffusion can occur in promoter search and define kinetic parameters that suggest facilitated diffusion occurs in promoter search even at the relatively high concentrations of RNAP σ^70^ holoenzyme in cells (0.2-0.6 μM; 12, 13).

## Results

### Real-time observations of single molecules of RNAP σ^70^ holoenzyme binding on DNA

We employed optical tweezers to hold single DNA molecules suspended between two optically trapped beads, such that the DNA and bound RNAP σ^70^ holoenzyme were completely free of surface interactions (13). Concurrent beam-scanning confocal microscopy (**Fig. 1A**) allowed us to visualize single molecules of Cy3B-labelled RNAP σ^70^ holoenzyme bound to the DNA in real-time (14). Representative consecutive confocal line-scans along the DNA were displayed as kymographs(14, 15), enabling analysis of binding dynamics (**Fig. 1B**). DNA binding events of Cy3B-labelled RNAP σ^70^ holoenzyme were visible as lines of elevated intensity above a background signal arising primarily from free Cy3B-labelled RNAP σ^70^ holoenzyme in solution (**Fig. 1B**). In these experiments, optical tweezers provided control of DNA tension. Most kymographs were acquired at 1 pN tension; some kymographs were acquired at 5 pN tension, yielding similar results (quantification below and in **SI**).

**Fig. 1.**
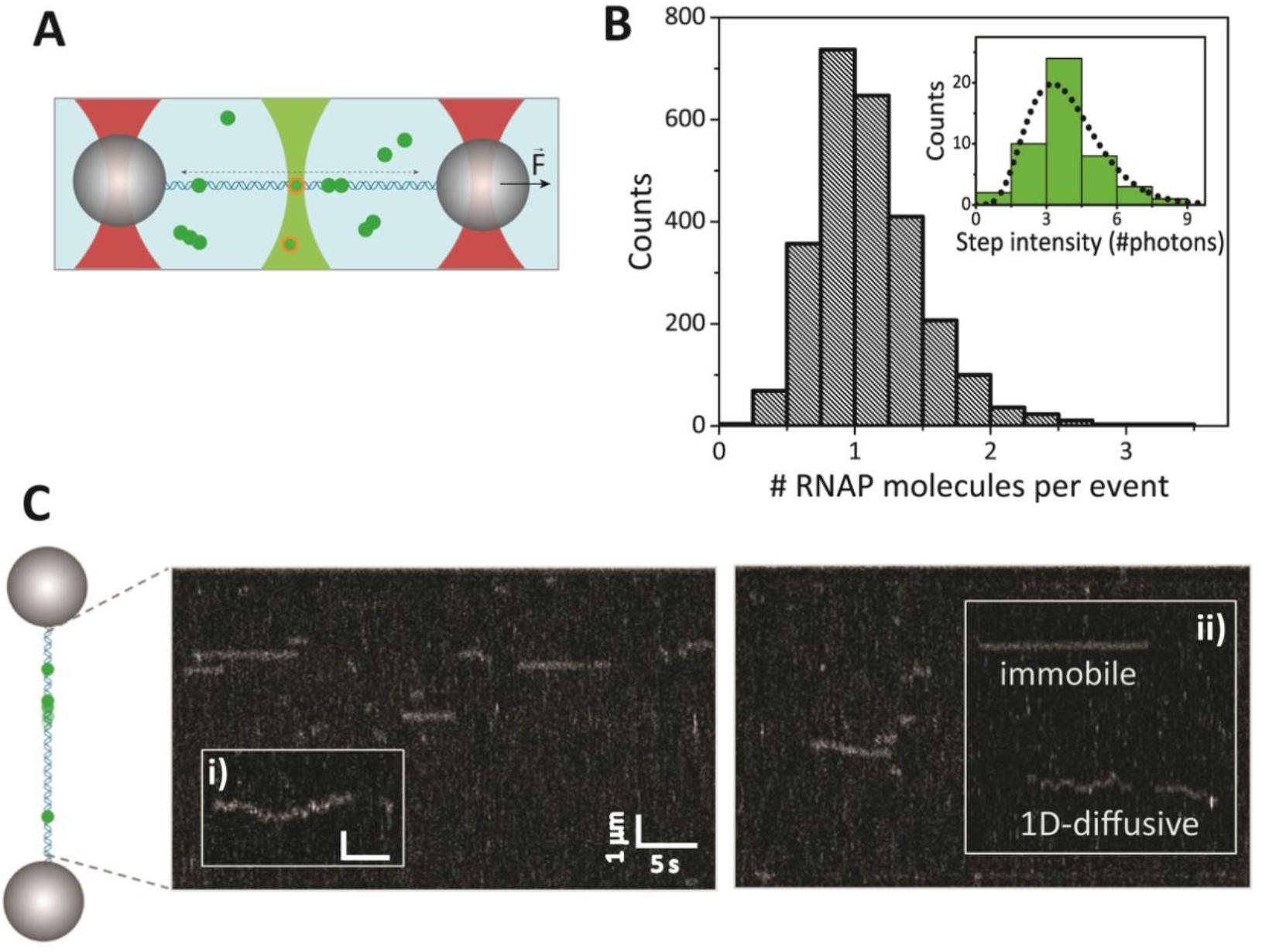
Single-molecule assay and experiments. (A) Schematic illustration of the assay showing a *λ*-phage DNA molecule tethered between two microbeads (grey), in the presence of Cy3B-labelled RNAP σ^70^ holoenzyme (green). A confocal excitation laser (light green) is scanned along the DNA to excite and image Cy3B-labelled RNAP σ^70^ holoenzyme, and two trapping lasers (red) trap the microbeads. (B) Distribution of initial peak intensities of Gaussian fits to confocal fluorescence line-scans across RNAP binding events detected at 1 pN DNA tension. Inset: distribution of single-step photobleaching intensities, used for calibrating single-molecule fluorescence intensities. Dotted line shows a gamma distribution fit to the data. (C) Bead-DNA-bead “dumbbell” (left) and kymograph, acquired by 1D scanning along the DNA, of single RNAP binding events (right; scale bar: 1μm x 5s). Inset i: example of a single 1D-diffusive RNAP binding event. Inset ii: example of a single immobile and a 1D-diffusive RNAP binding event. Scale bar: 0.6μm x 2s.

Kymographs were processed through single-molecule tracking to acquire information on the intensity, duration, location, and mobility of the fluorescently labeled RNAP (14, 15). The imaging parameters, including excitation intensity and line-scan rate, were optimized for a time-resolution of 90 ms. At these imaging settings, the trajectories exhibit single-step bleaching, a hallmark of single-molecule detection. The average photobleaching time of DNA-bound Cy3B-labelled RNAP σ^70^ holoenzyme was found to be 15 s (**SI**). All results were corrected for photobleaching. Photobleaching furthermore allowed calibration of the peak intensity of a single-dye (3.3±0.5 photons, see **Fig. 1B** and **SI**). Calibrated fluorescence intensities were used to assess binding stoichiometry, allowing the conclusion that the majority of observed DNA binding events represent single molecules of RNAP σ^70^ holoenzyme (**Fig. 1B**). A minority of events was excluded from further analysis because their fluorescence intensity was indicative of multimeric or aggregated RNAP (*N*_excluded_ =134 events of *N*_tot_ = 2600).

A first inspection of kymographs revealed non-diffusive RNAP binding events (“immobile events”), as well as 1D-diffusive RNAP binding events (**Fig. 1C**, inset i and ii). Extensive 1D-sliding was observed (**Fig. 1C**) and was found to be even more pronounced at lower Mg^2+^ concentration **Fig, S3**). 1D-sliding is further discussed below (see section “RNAP performs long-range 1D-sliding on DNA”). The observed 1D sliding did not correspond to transcription elongation, since the 1D sliding root mean-square displacement greatly exceeded that of transcription elongation under identical imaging conditions (**Fig. S4; SI**).

### RNAP binding events show sequence-dependent kinetics

By correlating RNAP binding and RNAP unbinding rates to locations along DNA, we confirmed previous reports (5, 6, 16) that RNAP DNA binding and unbinding kinetics are sequence-dependent. The frequency of observed DNA binding locations exhibited a characteristic asymmetric binding pattern along the analyzed bacteriophage-λ, DNA (**Fig. 2A**), correlating with the A/T-content along the DNA (red line in **Fig. 2A**).

**Fig. 2.**
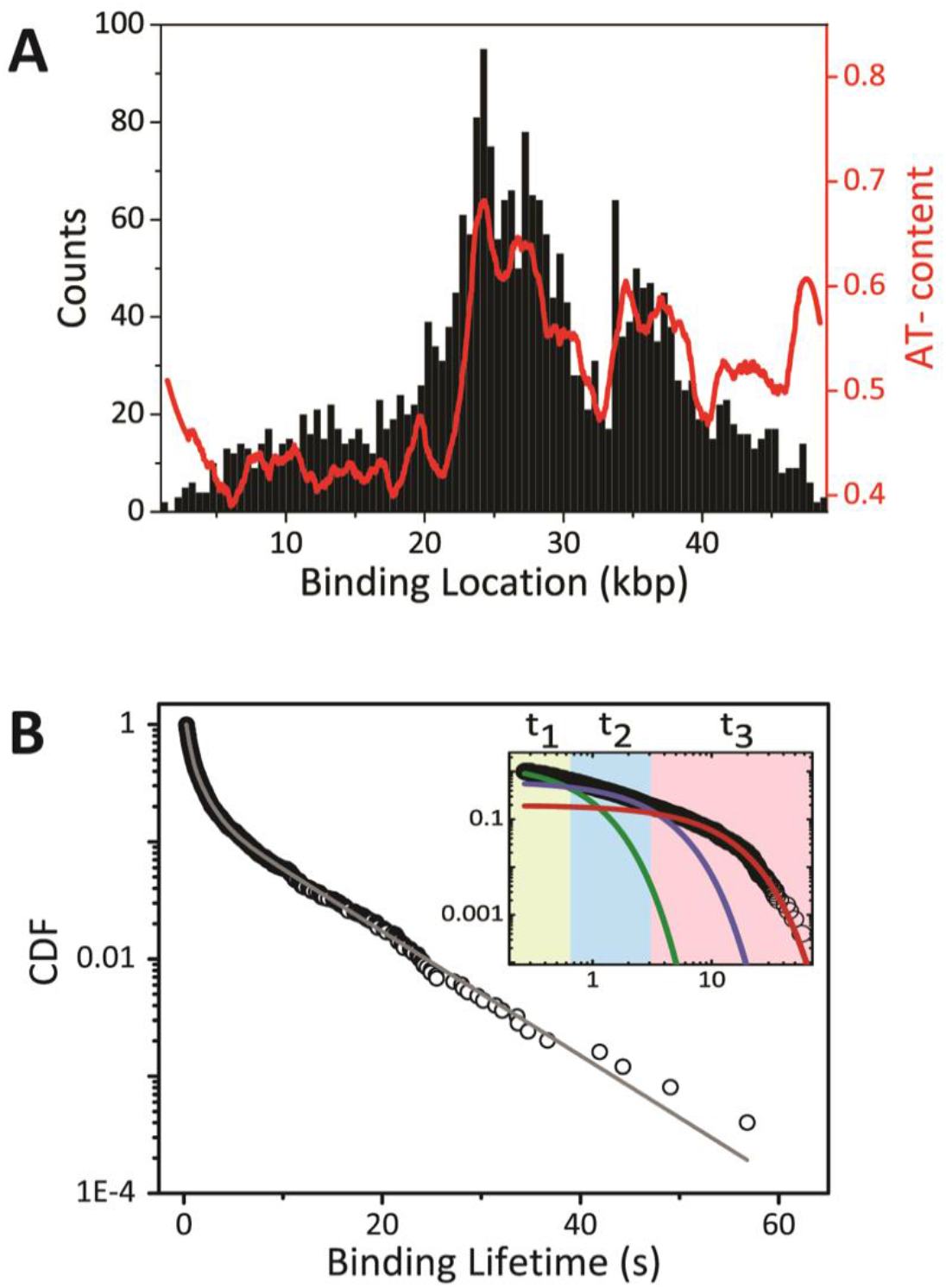
Binding locations and binding kinetics. (A) Histogram of locations of binding events along 48.5 kb *λ*-phage DNA, measured at 1 pN (N = 2466). The number of observed RNAP binding events correlates with local A/T-content (red line). (B) Normalized cumulative distribution function (CDF) of binding lifetimes of DNA binding events, measured at 1 pN (N = 2466). The solid line is the fit of triple-exponential function (see **Methods**). The inset shows the CDF data and the three single-exponential functions (green, blue and red line corresponding to populations with average lifetimes t_1_, t_2_, and t_3_, respectively) that make up the corresponding triple-exponential fit.

For the subsequent analysis of binding lifetimes, we imposed a minimum event duration 0.27 s (corresponding to 3 lines of 90 ms duration; see **Methods**) in order to minimize the chance of false-positive event detections in noisy kymographs. The cumulative distribution of event durations (**Fig. 2B**) could not be fitted with single-or double-exponential decay functions, but could be fitted satisfactorily with a triple-exponential function (**Fig. 2B** inset), indicating there are three distinct subpopulations of binding events, each with a characteristic lifetime and association rate (**Table 1**). At 1 pN, we distinguished a short-lifetime subpopulation (t_1_ = 0.37 s) comprising the majority, 53±3%, of binding events, an intermediate-lifetime subpopulation (t_2_ = 1.55 s) comprising 34±3% of binding events, and a long-lifetime subpopulation (t_3_ = 22.5 s) comprising 13±6% of binding events. Since t_3_ was considerably longer than the mean photobleaching time (22.5 s vs. 15 s), t_3_ was deemed a lower bound on the actual lifetime of the long-lifetime subpopulation (i.e., ≥22.5 s; see below). The characteristic event lifetimes (i.e., *t*_1_, *t*_2_, *t*_3_) measured at 5 pN DNA tension were directly comparable to those measured at 1 pN (**Table 1** and **Fig. S6**).

**Table 1.** DNA binding parameters obtained from analysis of RNAP binding events at DNA tension of 1 pN and 5 pN. Association rates are corrected to include missing events with lifetimes shorter than the *t_min_* (see **Methods**).

| | Dissociation Time (s) | | | | Association Rate ($M^{-1}s^{-1}$)$\cdot 10^8$ | | | Sliding parameters | | |
| --- | --- | --- | --- | --- | --- | --- | --- | --- | --- | --- |
| | $t_{1\_short}$ | $t_{2\_intermediate}$ | $t_{3\_long}$ | $t_3^\dagger$ | $k_{on\_1}$ | $k_{on\_2}$ | $k_{on\_3}$ | $t$ (s) | $D$ ($m^2/s$)$\cdot 10^{-14}$ | <distance> (kbp) |
| **1 pN** | $0.37 \pm 0.01$ | $1.55 \pm 0.05$ | $22.5 \pm 1.2^*$ | $300 \pm 10^{**}$ | $1.2 \pm 0.1$ | $0.8 \pm 0.1$ | $0.4 \pm 0.1$ | $1.9 \pm 0.3$ | $4.7 \pm 0.1$ | $1.2 \pm 0.1$ |
| **5 pN** | $0.22 \pm 0.01$ | $1.42 \pm 0.04$ | $23.2 \pm 2.9^*$ | | $4.6 \pm 0.1$ | $0.9 \pm 0.1$ | $0.2 \pm 0.1$ | $1.4 \pm 0.4$ | $4.5 \pm 0.3$ | $1.0 \pm 0.1$ |

\* Because the value of $t_{3\_long}$ exceeds the photobleaching time of 15 s, $t_{3\_long}$ should be considered a lower bound on the actual lifetime of the long-lifetime subpopulation; $^\dagger$ data acquired at low time resolution to obtain a reduced bleaching rate and thus a longer dissociation time (less limited by photobleaching). \*\* Since the value of $t_3^\dagger$ is not corrected for photobleaching, $t_3^\dagger$ should be considered as a lower bound on the actual lifetime of the long-lifetime subpopulation.

Finally, we analyzed association rates per DNA molecule using event frequencies. The average association rates (corrected for missed events, **SI**) at 1 pN for t_1_, t_2_, and t_3_ events at 1 pN tension were, respectively, 1.2· 10^8^ M^-1^s^-1^, 0.8· 10^8^ M^-1^s^-1^, and 0.4· 10^8^ M^-1^s^-1^, in line with observations by Harada et al. (5). Slight differences between association rates at 1 pN and 5 pN were observed that likely are attributable to tension-dependent kinetics and/or a tension-dependent signal-to-noise ratio (see further discussion below).

### RNAP binding lifetimes correlate with binding locations

To elucidate the identities of the three subpopulations of binding events, we inspected potential correlations in the spatial and temporal information of DNA-binding events. To this end, we divided events into three groups based on binding lifetimes (short, t ≤ 0.60 s; intermediate, 0.61 ≤ t ≤3.0 s; and long, t ≥ 3.1 s), according to the percentage of events (%) belonging to each subpopulation given by single-exponential fits, and mapped out binding locations along the DNA (**Fig. 3A**).

**Fig. 3.**
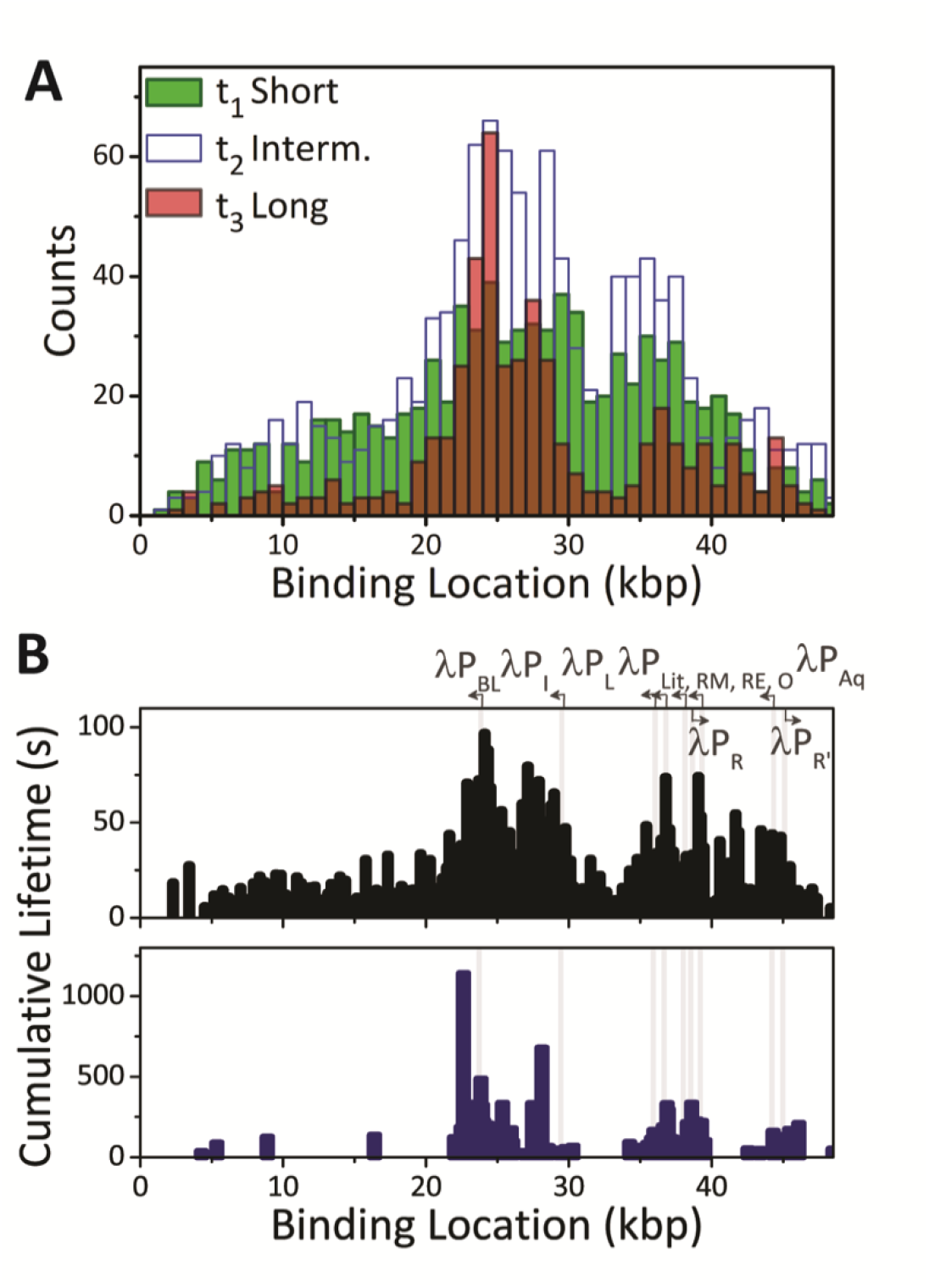
Position-dependence of binding kinetics. (A) Histogram of numbers of binding events as a function of binding positions along *λ* DNA, for three groups of event duration: short (t ≤ 0.60 s, green), intermediate 0.61 ≤t ≤ 3.0 s (blue), and long (t ≥ 3.1 s, red). (B) Cumulative binding lifetime as a function of binding location along *λ* DNA. Top plot: data recorded at high-temporal-resolution, high-photobleaching-rate conditions (black). Bottom plot: data recorded at lower-temporal-resolution, lower-photobleaching-rate conditions (dark blue). Arrows and vertical lines indicate locations of the 10 known RNAP-σ^70^-dependent promoters in *λ* DNA (17–21).

Although a complete separation of the three underlying subpopulations based on event duration was impossible, due to overlap in the exponential-lifetime distributions of subpopulations, clear trends were observed. The binding pattern of the long and intermediate events showed strong correlation with the local A/T-content of the DNA. In contrast, the binding pattern of the short events showed little or no correlation with the local A/T-content, indicative of non-specific binding. The most inhomogeneous binding pattern was observed for the longest-lifetime events. We propose that the longest-lifetime events are related to specific binding to promoter sites, which are located in regions of high local A/T-content (**Fig. 3B**). This proposal is supported by the observation that the cumulative binding time per location for all events exhibits peaks that correlated with the ten known RNAP-σ^70^-dependent promoter sites on *λ* DNA (17–21) (**Fig. 3B**, upper plot). The longest-lifetime events were limited by the rate of photobleaching. Accordingly, a second set of experiments was performed at lower time resolution in order to obtain slower bleaching rates. In this second set of experiments., the cumulative binding time for the longest-lifetime events correlated even more strongly with the ten known RNAP-σ^70^-dependent promoter sites on *λ* DNA (**Fig. 3B**, bottom plot).

### RNAP performs long-range 1D sliding on DNA

We further analyzed observed diffusive events to gain insight into potential contributions of 1D sliding to promoter binding (see example in **Fig. 1B**). A threshold event duration of t_cutoff_ (see Methods) was invoked to ensure that the average displacement due to 1D diffusion within this time was sufficiently large to be attributed to true motion along the DNA, rather than to artefacts of position errors, such as longitudinal thermal fluctuations of the DNA (~34 nm RMS at 1 pN) and experimental localization precision (~30 nm RMS at 250 nm FWHM diffraction-limited resolution and average of 13 photons per localization). With this threshold event duration, the 1D-diffusive events were found to constitute 15.5% of events longer than t_cutoff_ (N_Diff_/Nt>t_cutoff_ = 133/860) and 5.1% of all events. The average dwell time calculated from a single-exponential fit to the cumulative lifetime distribution of 1D-diffusive events with t > t_cutoff_ was 1.8 s (**Fig. 4A**, upper plot). The corresponding average 1D-sliding range was 400 nm, which equates to 1.2 kbp (**Fig. 4A**, bottom plot). Performing a mean square displacement analysis (15), we obtain a 1D diffusion constant for DNA-bound RNAP of 4.7· 10^-14^ m^2^/s, independent of DNA tension (**Fig. 4B** and **Table 1**). This value is as expected for rotation-coupled diffusion (22) of RNAP along the helical structure of DNA (~10^-13^ m^2^/s). Interestingly, the experimentally determined 1D diffusion constant suggests that RNAP experiences a relatively large, ~2*k_b_T*, average roughness in its sliding energy landscape (**SI**, further discussion below).

**Fig. 4.**
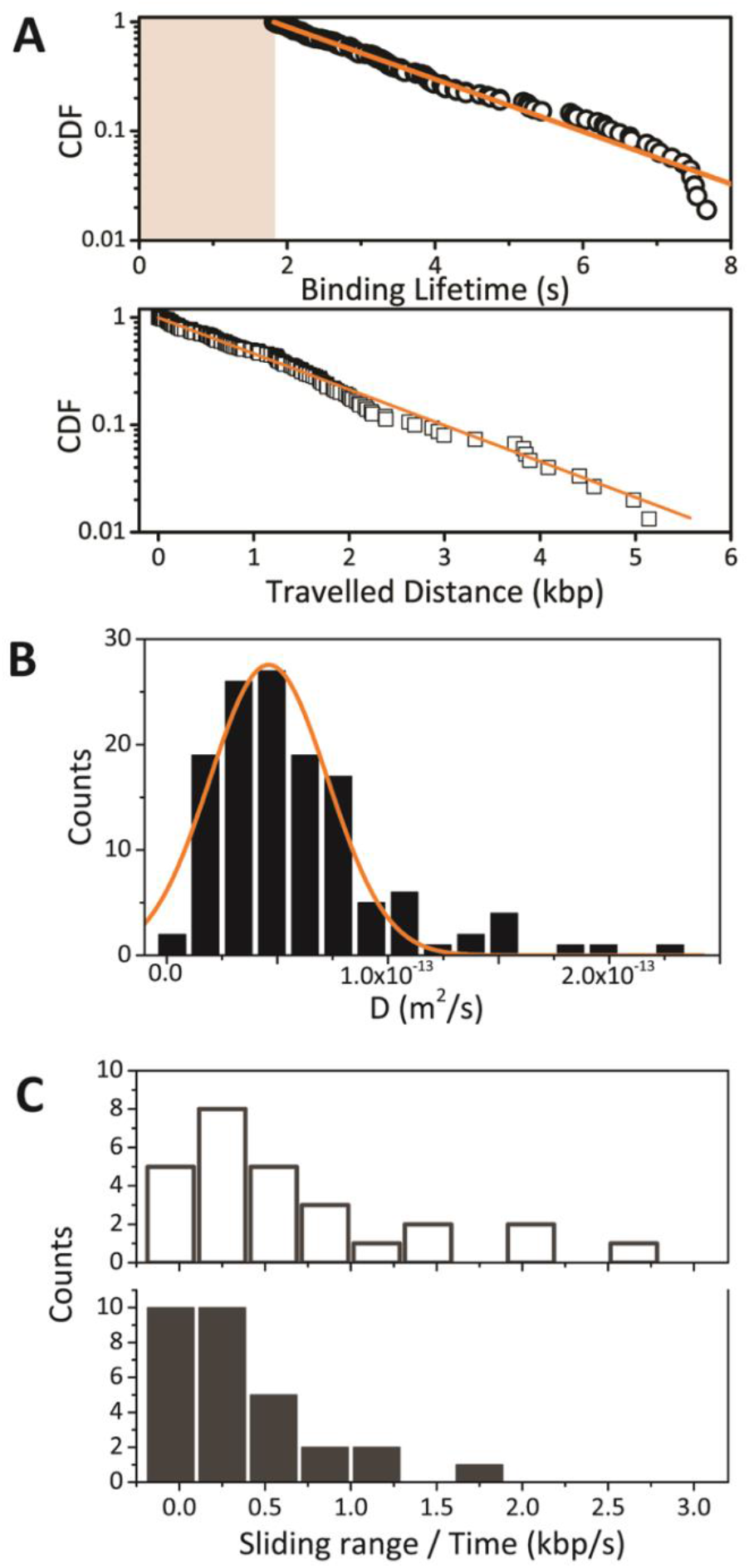
Analysis of RNAP 1D sliding. (A) Cumulative distribution function and single-exponential fit of binding lifetimes and correspondent travelled distance (kbp) of subset of sliding events with t > t_cutoff_, as indicated by grey background. (B) Histogram of diffusion constants obtained from mean-square-displacement analysis and corresponding Gaussian fit of individual tracked events with t > t_cutoff_. (C) Histograms of sliding range per unit time calculated for sliding events with t > t_cutoff_ that have start and end positions in promoter-free (top, white) or promoter containing regions of *λ* DNA (bottom, dark grey).

### Facilitated promoter search

Having observed that RNAP undergoes long-range 1D-sliding along the DNA, we sought evidence for facilitated diffusion – i.e., 1D sliding followed temporally by promoter binding. Unfortunately, unambiguous direct monitoring of individual facilitated diffusion events was not possible, due to noise in the position data and the stochastic nature of 1D-diffusion. As an alternative to direct monitoring of individual facilitated diffusion events, we compared mean 1D-sliding distances for promoter-containing vs. promoter-free regions of the phage *λ* genome. We made use of the fact that phage *λ* genome contains multiple micrometers-long stretches of DNA devoid of promoters (**Fig. 3B**). We reasoned that, in promoter-free regions, mean 1D-sliding distances should be unperturbed by promoter binding, whereas, in promoter-containing regions, mean 1D-sliding distances should be reduced by promoter binding. We find that the mean sliding distances, corrected for event durations, are >2-fold greater in promoter-containing regions (0.12±0.06 kb/s) than in promoter-free regions (0.27±0.05 kbp/s) (**Fig. 4C**), indicating that facilitated diffusion contributes to promoter binding under the conditions analyzed.

## Discussion

We investigated the promoter-search dynamics of RNAP σ^70^ holoenzyme in real time at the single-molecule level. Our data confirm previous findings that RNAP σ^70^ holoenzyme exhibits sequence-dependent DNA binding kinetics (5, 6). Binding events could be divided into short non-specific binding events, longer binding events on A/T-rich DNA, and longest binding events on promoter DNA, in line with previous results (5, 6). Imaging with lower temporal resolution in order to obtain lower photobleaching rates, we showed that the longest events have lifetimes >100-1000 s and occur at promoter sites, characteristic of RNAP-promoter transcription initiation followed by transcription elongation complexes. Indeed, directed motion characteristic of transcription elongation can be observed for such long events (**SI**).

DNA tension is found to affect RNAP association rates, rather than the dissociation rates. More specifically, at 5 pN tension, we observe an apparent elevation of the association rate for nonspecific DNA and a slight reduction of the association rate for promoter DNA, compared to binding at 1 pN. The latter result confirms the tension-dependent reduction of promoter binding observed by Harada et al. (5) and is reminiscent of the observed tension-dependence of promoter binding for the structurally unrelated bacteriophage T7 RNA polymerase observed by Skinner et al. (23). We attribute the apparent elevated association rate for short lifetime events at non-specific DNA to enhanced signal-to-noise ratios at high tension, due to suppressed motions of DNA molecules resulting in reduced motion blurring of fluorescence images (14, 24).

A striking feature of our data is the demonstration of long-range 1D sliding along DNA by bound RNAP σ^70^ holoenzyme (**Figs. 1C, 4**). We are confident these sliding events involve single molecules of RNAP σ^70^ holoenzyme, because we excluded data from aggregates of RNAP σ^70^ holoenzyme based on the calibrated fluorescence intensities (see **SI**). The average sliding time in our data is 1.8 s, and the average covering distance travelled is 1.0-1.2 kb.

Mean-square-displacement analysis yielded a 1D diffusion constant of 4.7· 10^-14^ m^2^/s, in excellent agreement with findings of Harada et al. (5; estimated to be on the order of 10^-14^ m^2^/s) and in agreement with expectation for rotation-coupled 1D diffusion of RNAP around the helical pitch and along the helix axis of DNA (23; 10^-14^ m^2^/s). Our analysis of 1D sliding events was restricted to events longer than t_cutoff_= 1.9 s. It seems likely that shorter events, which were excluded in our sliding analysis by the cutoff time, also exhibit 1D sliding through rotation-coupled diffusion. Assuming the same diffusion constant, D, of 4.7· 10^-14^ m^2^/s, the average lifetime of short-lifetime binding events (0.3 s) would imply an average sliding range of 500 bp for short-lifetime binding events. This sliding range is consistent with sliding ranges of 300 bp and 440 bp reported in early studies (4, 5).

Our observation of unambiguous 1D-sliding by RNAP σ^70^ holoenzyme contrasts sharply with results of Wang et al.(6), who failed to detect 1D-sliding in a surface-tethered DNA-curtain assay, even though their range of salt conditions overlapped ours. We tentatively attribute their failure to detect 1D sliding to their use of an experimental design that imposed two sets of non-native constraints on 1D sliding, particularly rotation-coupled 1D sliding (22): (i) non-native hydrodynamic drag by the nanoparticle used to label RNAP in their experiments [diameter = 26 nm (6); ~2x diameter of RNAP (25)], and (ii) non-native surface interactions by DNA, RNAP, and the RNAP-conjugated nanoparticle within the surface-tethered DNA curtains used in their experiments [mean distance between DNA and surface = 15 nm (6); ~1x diameter of RNAP and ~0.6x diameter of RNAP-conjugated nanoparticle]. Wang et al. (6), employed an experimental design that magnified the hydrodynamic drag on RNAP and that required RNAP and an RNAP-conjugated nanoparticle to thread through a gap between DNA and surface that averaged just ~1x the diameter of RNAP and just ~0.6x the diameter of the RNAP-coupled nanoparticle in each 10 bp of rotation-coupled 1D sliding. Further evidence for non-native surface interactions in Wang et al. (6) is found in the lifetime of the shortest binding events observed by Wang et al (0.03 s), which is <1/10 the lifetime of the shortest binding events observed in the absence of surface interactions (0.37 s; **Table 1**).

What can we finally conclude regarding the relevance of facilitated diffusion for promoter search? Does 1D sliding contribute to promoter binding? Our data are consistent with 1D-sliding contributing to promoter binding. A theoretical model of facilitated diffusion of Slutsky (26) suggests that proteins that undergo facilitated diffusion exhibit at least two DNA binding modes: a search mode with weak protein-DNA interaction (~1-2 k_B_T), and a recognition mode at the specific site with a protein-DNA interaction energy (> 5 k_B_T). Based on analysis of our data in terms of rotation-coupled 1D diffusion, we estimate an interaction energy difference between 1D-sliding and long events that is indeed ≥ 3 k_B_T higher for a recognition mode (**SI**), in agreement with Slutsky (26). Interestingly, the observed average roughness of the sliding energy landscape, ~2 kBT (**SI**), is relatively high compared to roughness values found for other DNA-binding proteins by Blainey et al. (between 0.75 and 1.35 k_B_T; 23). We attribute this relatively high roughness value to the unusually large length of the DNA segment sequence-specifically contacted by RNAP σ^70^ holoenzyme (up to ~120 bp; 28) and/or to the unusually large number of distinct, functionally independent, sequence-specific DNA binding modules present in RNAP σ^10^ holoenzyme (up to six distinct, functionally independent, sequence-specific DNA binding modules: αCTD^I^, αCTD^II^, σ^70^ region 4, σ^70^ region 3, σ^70^ region 2, σ^70^ region 1.2, and β pocket; 28).

Finally, our data on DNA binding kinetics allows us to estimate the “facilitation threshold concentration” C_0_, the RNAP concentration at which the rate of promoter binding due to 3D diffusion equals that due to facilitated 1D diffusion. To obtain a lower bound for the facilitation threshold, we use the intermediate event lifetime τ = t_2_ and find that C_0_ ≥ 0.2 μM (see **SI**). Strikingly, this facilitation threshold concentration is comparable to, or even larger than, the intracellular concentration of RNAP σ^70^ holoenzyme in *E. coli* of 0.2-0.6 μM (11, 12). Thus our results suggest that facilitated 1D diffusion plays significant roles at RNAP σ^70^ holoenzyme concentrations in cells. It should be noted that our *in vitro* system is drastically simplified compared to the *in vivo* situation, where molecular crowding, roadblock proteins on DNA, and other factors are likely to affect both 3D diffusion and facilitated 1D diffusion.

Nevertheless, the finding that the facilitation threshold concentration equals or exceeds the concentration of RNAP σ^70^ holoenzyme *in vivo* makes it tempting to speculate on effects of facilitated 1D diffusion on transcription rates *in vivo*, and, in particular, tempting to speculate that stochastic or environment-induced fluctuations in concentrations of RNAP σ^70^ holoenzyme *in vivo* could be “buffered” around the facilitation threshold concentration, with concentrations of RNAP σ^70^ holoenzyme above the threshold resulting in lower efficiencies of promoter search, and concentrations of RNAP σ^70^ holoenzyme below the threshold resulting in higher efficiencies of promoter search.

## Methods

### Proteins, DNA, and buffers

For experiments in Figs. 1-4, S1-S2, and S4-S6, Cy3B-labeled *E. coli* RNAP σ^70^ holoenzyme (Cy3B incorporated at position 284 of RNAP β’ subunit; labeling specificity >90%; labeling efficiency ~90%) was prepared by unnatural-amino-acid mutagenesis of a gene encoding RNAP β’ subunit to incorporate 4-azidophenylalanine at position 284 of β’ as described(28, 29), followed by azide-specific Staudinger ligation to incorporate Cy3B at position 284 of the resulting β’ derivative as described (28, 29), followed by core-enzyme reconstitution using the resulting Cy3B-labelled β’ derivative and RNAP β, FLAG-αNTD^I^-αNTD^II^ and rø subunits as described (28, 29), followed by holoenzyme reconstitution by combining the resulting Cy3B-labeled core enzyme derivative and 4 mole equivalents of σ^70^ in imaging buffer (20 mM Tris-HCl pH, 50 mM KCl, 4 mM MgCl2, 1 mM dithiothreitol, 1 mM ATP, 1 mM CTP, 1 mM UTP, and 0.25 mM GTP) for 30 min at 30°C. For experiments in Fig. S3, Cy3B-labeled *E. coli* RNAP σ^70^ holoenzyme (same labeling site, same labeling specificity, and same labeling efficiency) was prepared in the same manner, but replacing FLAG-αNTD^I^-αNTD^II^ by FLAG-αNTD^I^-αNTD^II^-HaloTag [prepared in the same manner as FLAG-αNTD^I^-αNTD^II^, but using plasmid pET28a-FLAG-αNTD^I^-αNTD^II^-HaloTag, constructed from plasmid pET28a-FLAG-αNTD^I^-αNTD^II^ by inserting a SpeI site (ACTAGT) and a HaloTag coding sequence (nucleotides 169-1059 of plasmid pH6HTC His6HaloTag T7 vector; Promega, Inc.; Genbank JN874647) immediately upstream of the stop codon of the FLAG-αNTD^I^-αNTD^II^ coding-sequence. Bacteriophage *λ* DNA, 48.5-kbp long, was biotinylated as described previously (30). Streptavidin-coated polystyrene beads (diameter 1.9 μm, SpheroThec) were optically trapped in a flow of PBS buffer inside a multichannel laminar-flow system. By moving the flow cell perpendicular to the flow direction, trapped beads are subsequently immersed in an adjacent flow lane containing *λ*-DNA in PBS, where single DNA molecules are tethered between the two beads. The DNA is mechanically characterized in imaging buffer containing 0 or 5 nM Cy3B-labeled RNAP σ^70^ holoenzyme at 23°C.

### Optical trapping and fluorescence microscopy

the experiments were performed on a custom-built instrument that combines dual-trap optical tweezers with beam-scanning confocal fluorescence microscopy, as described previously (14).

### Single-molecule force-fluorescence assay

DNA-bound Cy3B-labeled RNAP was imaged by beam-scanning confocal microscopy as described previously (14). Kymographs are acquired by scanning a 542 nm excitation laser spot (86 μW before objective) along the optically stretched DNA with a pixel dwell time of 0.02 ms, pixel size 75 nm, and line-to-line time of 30 ms. In the bleaching study the line-to-line time was varied between 30ms – 90ms. See also SI.

### Kymographs Time-Spatial resolution (e.g. summing lines in analysis)

In order to reduce fluctuations in the background signal associated primarily with proteins that are freely diffusing in solution, we sum the photon counts acquired at each position over three subsequent line scans resulting in a reduced temporal resolution of 90ms. Furthermore, binding events shorter than three of such summed lines (i.e. t < 270 ms) are excluded from analysis.

### Determination of minimum event duration for analyzing 1D diffusion

A first estimate of the diffusion constant was obtained from an initial subset of diffusive events with event duration longer than 3s. This minimum event duration was chosen because over such long time interval, the displacements of the random walk of the RNAP are sufficient to unambiguously identify 1D-diffusive behavior from a visual inspection of the kymographs. Based on the mean-squared displacement of this subset of diffusive traces, we calculated the diffusion constant 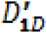 as explained previously(15). Based on this first estimate of the diffusion constant, we then refined the event duration threshold for analyzing diffusive traces by calculating the time after which the average displacement of a random walk with 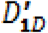 equals the typical point-spread function of our confocal microscope of 300 nm, yielding a refined cutoff time of 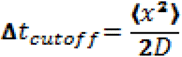, where x is the size of our point spread function (~300 nm). This yields 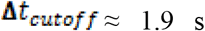. Finally we selected only diffusive traces, for which 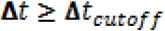.

### Correction of diffusion constant for changes in relative DNA extension

The diffusion constant directly extracted from the MSD analysis *D_MSD_* were corrected for the end-to-end distance *L_B_* of the DNA at the two different forces used: at 1 pN is 13.3 μm and at 5 pN is 15.4 μm. The corrected diffusion constant is 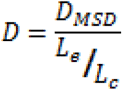. where *L_C_* is the *λ*-phagc DNA contour length (16.49 μm).

## Supporting information

## Acknowledgement

This work was supported by VICI grants (G.J.L.W. & E.J.G.P.), VIDI grants (W.H.R. & I.H.), a VENI grant (I.H.) from the Nederlandse Organisatie voor Wetenschappelijk Onderzoek, a European Research Council starting grant (G.J.L.W.), and grant GM041376 from the National Institutes of Health (R.H.E).

## Author contributions

I.H., G.J.L.W., E.J.G.P. and R.H.E. conceived the study. I.H. designed the experiments. M.M., and A.Mal., performed single-molecule experiments and analyzed data. A.Maz., A.C. and R.H.E. provided labeled proteins. I.H. built the used single-molecule instrumentation. I.H., G.J.L.W., E.J.G.P., W. H. R. and R.H.E. advised on experiments and data interpretation. I.H., M.M., G.J.L.W. and R. H. E. wrote the manuscript. I.H. and G.J.L.W. lead the research.

