## Supplementary material for "One-dimensional sliding assists σ^70^-dependent promoter binding by Escherichia coli RNA polymerase"

**This PDF file includes:**

Supplementary text
Figs. S1 to S6
References for SI reference citations

### SI Materials and Methods

**Intensity calibration.** For analyzing the DNA-binding stoichiometry, the fluorescence intensity collected from one dye (intensity calibration) was estimated by quantifying the bleaching steps exhibited by multi-dyes events, which yielded a peak intensity (amplitude of a Gamma distribution fit to the intensity profile) of 3.3±0.5 photons per dye, see Fig 1B inset, in the main text. This value of the intensity of a single dye is compared to the initial intensity of the observed DNA-binding events, which is plotted in a histogram (Fig. 1B). The initial intensity exhibits a single peak of average intensity of 3.6±0.3 photons, in excellent agreement with the intensity of a single dye. We conclude that no oligomers of RNAP are present in our analyzed data. To nevertheless exclude rare RNAP multimers from our analysis, we only used traces with an initial intensity within two standard deviations obtained from the fit.

**Bleaching study.** In this study we identify DNA-bound proteins using single-molecule fluorescence. Association and dissociation of proteins can be identified through analysis of trajectories with elevated fluorescence intensity in a kymograph. However, the ending of such a trajectory can relate to dissociation, but also to photo-bleaching of the fluorophore. For a correct analysis, we first quantified after how much time, on average, the used dye Cy3B bleaches under our experimental conditions. To this end we studied the DNA-interaction lifetimes of our labeled-RNAP for three different values of the line-to-line time $t_F$ (30 ms, 60 ms, 90 ms). We expect to observe longer traces for a longer $t_F$, due to reduced fluorescence exposure. The observed time $t_0$ (calculated as the exponential decay time constant from a statistics on the interaction lifetimes) was calculated and analyzed as function of the corresponding line-to-line time. The resulting linear relation (Fig. S1A) allowed to distinguish between the observed time, the protein real interaction time, and the photo-bleaching time (as number of frames before photo-bleaching occurs). The latter value was found to be 490 lines, and it was used to correct all our experimental results on the interaction lifetimes.

**Determining the binding position on DNA.** The position of a protein along the DNA was calculated with a custom-made tracking analysis software. We extracted three values for each event location: initial position, final position and mean position, calculated respect to the beads position in each kymograph (i.e. used to correct for the error introduced by thermal motion and instrumental bead positions variation). A Gaussian fit of the beads centers, which emits with higher intensity, was used to estimate the positions of tethered DNA molecule. From this reference point, the position (in nm) of every protein bound on the DNA was obtained. We normalized their locations respect to the bead-to-bead distance (13.3 μm and

15.4 μm at 1 pN and 5 pN respectively) and we converted from nm to kilo-basepairs (kbp, where 1 bp = 0.34 nm). The used λ-phage DNA is 48.502 kbp-long.

Because the bead-DNA attachment is identical on both DNA ends, the orientation of the DNA is in principle random. For the position analysis, we first wanted to attempt to correct for this random orientation of our DNA molecules. The right DNA orientation is given the locations of stable traces, under the assumption that stable traces are proteins bound to promoter sites which are located on one side of the DNA (no promoter sequence is found on the other side). Moreover RNAP has a higher affinity for the AT-rich regions, leading to more observed binding events on one half of the λ-phage DNA molecule. These two considerations were used to orient each DNA molecule and obtain the relative protein positions on it (Fig. S1B).

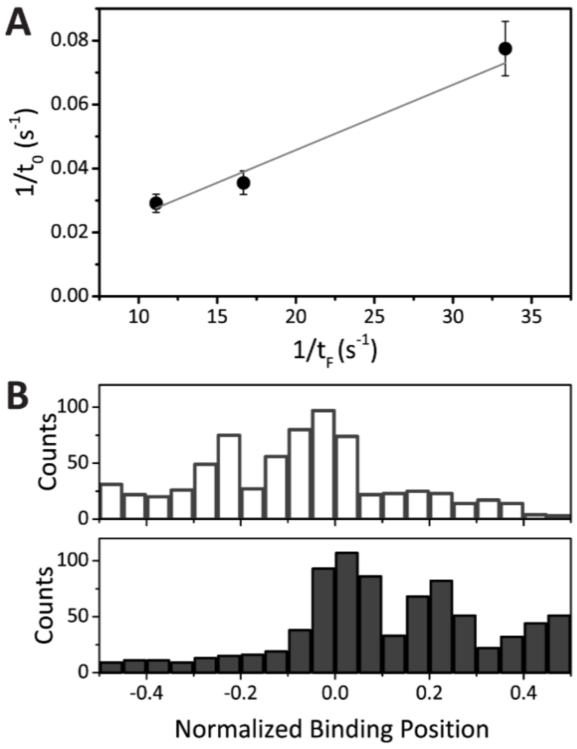


**Fig. S1** (A) Bleaching study: the observed binding lifetimes (y-axis) for three different inter-frame wait time (x-axis) results in a liner relation that gives an estimation of the number of scanned line before bleaching of a single dye occurs. The bleaching time is used to correct all measurements. (B) Two examples of binding events on two different DNA molecules imaged for several minutes. The clear signature of an higher affinity for one side of the DNA (i.e. rich in AT base pairs) allows to orient each DNA molecule before to unify all of them as one dataset.

**Kymograph resolution specifications.** A visual inspection of kymographs with the chosen setting does not allow to detect all diffusive traces, therefore the mean-square-displacement analysis is performed with an automated tracking software. More specifically: the chosen pixel size for the fluorescence confocal scanning is 75 nm which correspond to ~220 bp of DNA, while the confocal diffraction-limited spot is ~250 nm or ~735 bp. Any displacement that can be detected from visual inspection of the kymographs is of the order of the spot size, that correspond to a sliding range over hundreds to thousands of basepairs. Through particle tracking of the kymographs we substantially improve our ability to detect displacements: our localization accuracy (at about 10 photons per localization) is about 30 nm, corresponding to about 90 bp.

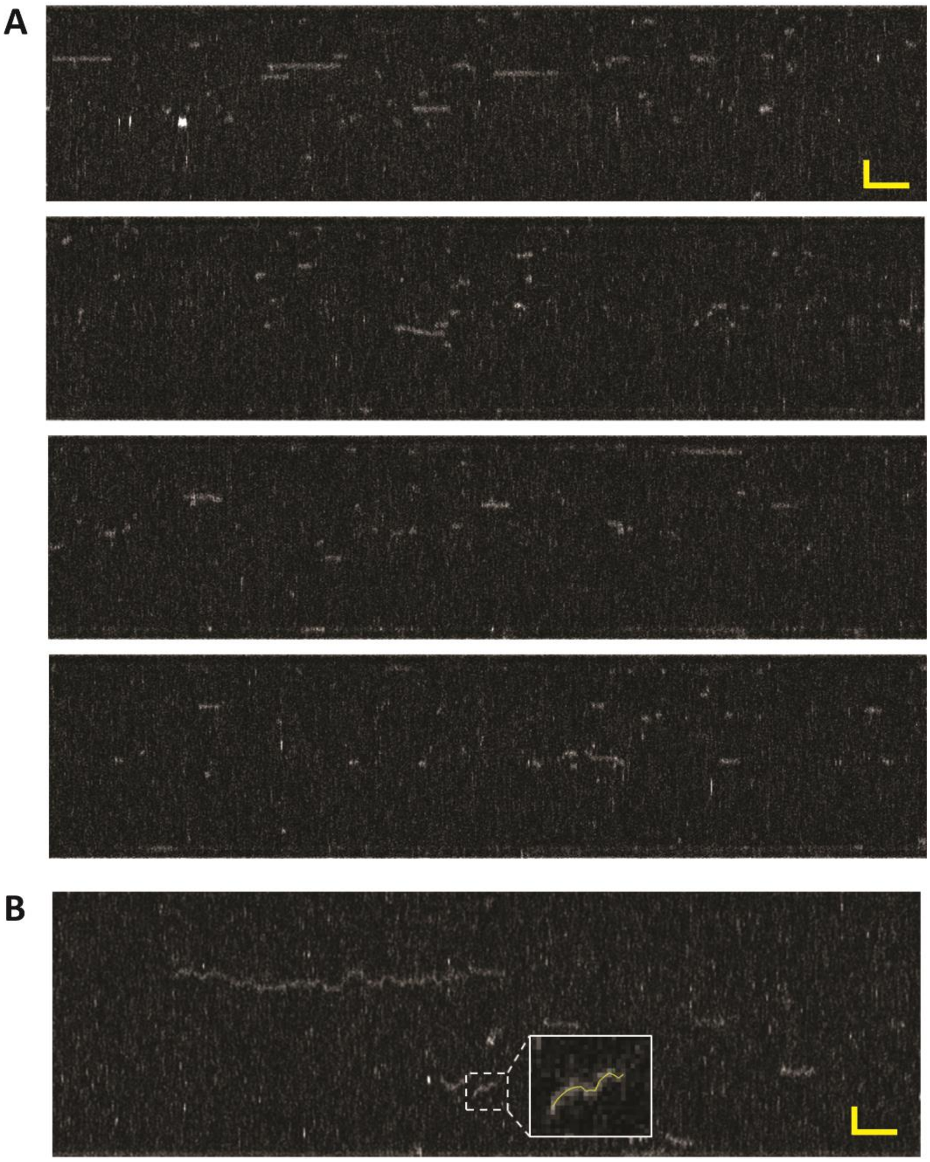


**Fig. S2**: (A) Examples of recorded full length kymographs. Scale bar: 2µm x 5 s. (B) Zoom into example of long-range sliding RNAP and shorter traces. Inset: zoom into one trace tracking example. Scale bar: 1.5µm x 2 s.

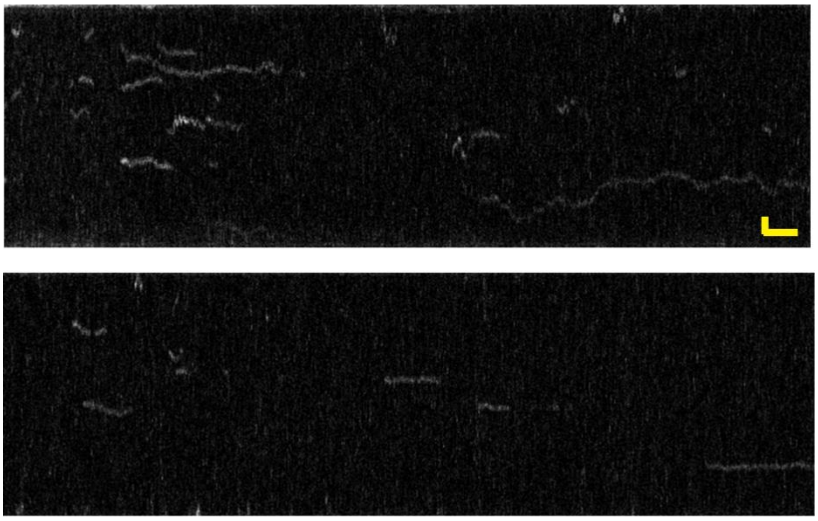

**Fig. S3**: Examples of kymographs recorded at lower salt conditions (20 mM Tris-HCl, 50 mM KCl, 1 mM $MgCl_2$, 1 mM DTT, ribonucleotides (rNTPs) 1 mM of ATP, CTP and UTP, and 0.25 mM of GTP). For illustrative purposes we have analyzed protein binding at 1 mM $MgCl_2$ (N=528) at 5 pN tension. We observe reduced event duration of short and intermediates events ($t_1$=0.16±0.02s, $t_2$=0.92±0.06s), while the long event duration was found to be exactly the same ($t_3$=22.4±1.1 s). Interestingly the frequency of diffusion events at lower salt conditions (1 mM $MgCl_2$) is 3-fold higher (18% of total events) and their durations is 3-fold longer (6.5±0.5 s), data not shown.

**Transcription traces.** Transcription cannot explain the apparent sliding we observe: under the experimental conditions, we expect a transcription rate of ~ 10 nt/s, which translates to a displacement of ~ 5 nm/s. For a trace length of 2 s, this would thus lead to only 10 nm displacement. Since 10nm is far below the diffraction limited spot size of 250 nm, the pixel size of 75 nm, the longitudinal thermal fluctuation of the DNA of 34 nm, and the localization accuracy of ~ 30 nm, transcription could not visually nor in analysis be distinguished from an immobilized RNAP, nor be mistaken for a diffusing one. For comparison, 1D-diffusion at the measured diffusion constant of $D=4.7\cdot10^{-14}$ $m^2/s$ corresponds, for a 2 s trace length, to an average absolute displacement of $\sqrt{(2Dt)}$ = ~500 nm or ~1500 bp. The visually striking examples are merely stochastically infrequent events with sufficiently long lifetime in the tail of the exponential lifetime distribution of the sliding events.

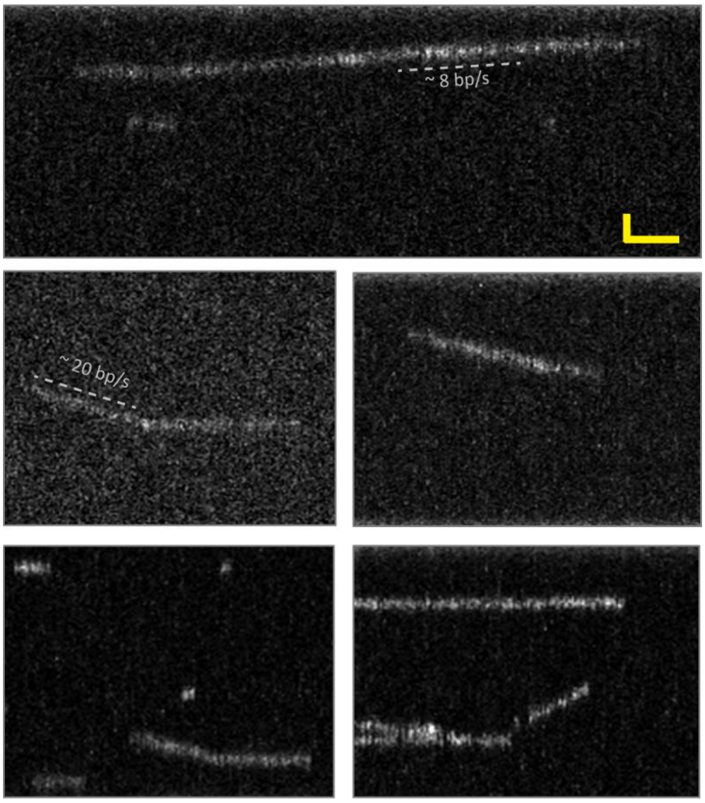


**Fig. S4**: Examples of transcription traces that could be observed with low bleaching fluorescence settings. Scale bar: 1 μm x 75 s

**Correction of cumulative event duration for missed events.** The cumulative distribution function of the binding lifetimes is fit as a three-exponential decay (namely the sum of three single exponential decays, see main text). This fit reveals how many events belong to each exponential decay and thus to each off-rate population. The contribution of each population to the total (including both observed and not-detectable) events are given by the exponential multiplication factor (i.e. the exponential function at time zero). This analysis takes into account also the missing events, or not-detectable because of our time-resolution limit ($t_{min}$), leading to the populations percentages showed in the main text. In order to correct for the missing events that contribute to the RNAP association rate we need to consider the intersection of the exponential function $t_{min}$ = 0.27s. To this end we multiply each association rate for the ratio between the exponential multiplication factors at 0 s and at 0.27 s.

**Rotational 1D-diffusion constant.** The observed diffusion constant was calculated under the assumption that RNAP undergoes 1D-diffusion along the DNA. We now compare this to the theoretical diffusion constant from the helical diffusion model (1): $D_{\text{helix slide}}^{\text{Theor.}} \approx b^2 \frac{k_B T}{14\pi\eta R^3} F(\varepsilon)$. Where b = 3.4 nm (10.5 bp) is the distance per full rotation around the DNA; $R$ is the protein radius (we assume that the distance between the DNA axis and the protein axis equals the protein radius); $\eta = 10^{-3}\,\text{Pa}\cdot\text{s}$ is the viscosity of water at about 20°C; $F(\varepsilon) = e^{-\left(\frac{\varepsilon}{k_B T}\right)^2}$ is a retarding factor due to the roughness of the energy landscape of a protein that moves along the DNA helix, where the average free-energy barrier for sliding along the DNA was calculated for many proteins by Blainey et al. (1) to be $\varepsilon \approx 1\,k_B T$. We estimate the radius of our Cy3B-labeled RNAP to be ~7.5 nm (this is an upper limit given by the sum of the radius of the holoenzyme to be ~7.0 nm(2) and of the Cy3B dye to be ~0.5 nm, obtained from its molecular weight. We obtain $D_{\text{helix slide}}^{\text{Theor.}}(\text{Cy3B} - \text{RNAP}) \approx 9 \cdot 10^{-13} \frac{m^2}{s}$. This theoretical value for the diffusion constant is higher than the value that we found from our experimental analysis: $D_{1D}^{\text{Observed}} < D_{\text{helix slide}}^{\text{Theor.}}$. A potential explanation for the discrepancy lies in the roughness of the energy landscape that was assumed to be $\approx 1\,k_B T$. Since RNAP is known to interact very strongly with the DNA (at least during transcription), we speculate that $1\,k_B T$ underestimates the true energy landscape between DNA and RNAP. Given the experimentally found value for the diffusion constant $D_{1D}^{\text{Observed}} = (4.7 \pm 0{,}2) \cdot 10^{-14} \frac{m^2}{s}$, we used the previous equation for $D_{\text{helix slide}}^{\text{Theor.}}$, to calculate the roughness of the energy landscape: $\varepsilon_{\text{RNAP}} \approx 2\,k_B T$. This suggests that RNAP interacts more strongly with the DNA compared to the mean value calculated by Blainey et al. for different sliding proteins (see main text for further discussion).

**Estimate of the energy difference between search mode and recognition mode** We estimated the difference in binding energy ΔG between the 1D-sliding mode (search mode) and the target-bound mode (recognition mode). To this end we calculated $\Delta G = k_B T\ ln(K_{D,search}/K_{D,recognition})$, where the dissociation constant $K_D = \frac{k_{off}}{k_{on}}$ is the ratio between the dissociation rate and the association rates. In order to estimate the dissociation constant of the search mode, we use the kinetic rates that we measured for short lifetime population (population 1: $k_{off} = 2.7\ s^{-1}$, $k_{on} = 1.2 \cdot 10^8\ M^{-1}s^{-1}$ , see Table 1). For a lower-limit estimate of the dissociation constant of the recognition mode, we use the kinetic rates that we measured for the long lifetime population (population 3: $k_{off} = 0.04\ s^{-1}$, $k_{on} = 0.4 \cdot 10^8\ M^{-1}s^{-1}$, see Table 1). Note that for the long lifetime population $k_{off}$ is limited by photobleaching, such that this value represents an upper limit. We thus find a lower limit estimate for the energy difference of ≥3 $k_BT$ between the search mode and recognition mode.

**Facilitation threshold estimation.** The facilitation threshold concentration $C_0$ is defined as the protein concentration at which the binding rate to find a target on the DNA through facilitated diffusion equals the binding rate to find that target through 3D diffusion (3). As described in the supplementary information of reference (3)(equation S6-S8 and supporting text), we can estimate a lower bound to the facilitation threshold concentration as $C_0 \geq \frac{1}{\tau k_{CC}}$, where τ is the binding lifetime for sequence nonspecific RNAP-DNA complexes, and $k_{CC}$ is the rate constant for forming the closed complex at the promoter site (3). We estimated $k_{CC} \leq 4 \cdot 10^6\ M^{-1}s^{-1}$ as described below (see section Estimation of the rate of closed complex formation). We thus find that $C_0$ ≥ 0.2-0.7 μM for τ ranging from nonspecific interaction times of short events ($t_1$ = 0.37s) to those of intermediate events ($t_2$ = 1.55s). This estimation shows that at our experimental concentration (5nM) and conditions, facilitated diffusion contributes to the promoter search process.

**Estimation of the rate of closed complex formation.** In order to find a lower limit of the facilitation threshold concentration according to the procedure described above, we estimate an upper limit for the rate of closed complex formation. To this end, we quantified the average binding rate under low bleaching conditions, where the minimum event duration that we analyzed was 3s with an average lifetime of 300s. Our assumption is that the majority of these events constitutes promoter binding events followed by closed complex formation. Indeed, Fig. 3 B in the main text shows that these events correlate strongly with known promoter sites. For each DNA molecule, we analyzed only the time to initial binding, i.e. the very first binding event longer than 3s, in order to ensure that all promoters are unoccupied. Fig. S5 shows the normalized cumulative distribution of initial binding rates at promoter sites. An exponential fit to this distribution yields $k_{\mathrm{CC}} = 4 \cdot 10^6\ M^{-1}s^{-1}$. We note that phage λ-DNA contains multiple promoter sites (as indicated in the main text) available for binding, which will increase the apparent rate of closed complex formation in our analysis. However, each promoter may exhibit different RNAP binding kinetics. In order to estimate a lower bound for the facilitation threshold concentration we used an upper limit for the rate of closed complex formation and thus do not correct for the number of promoter sites.

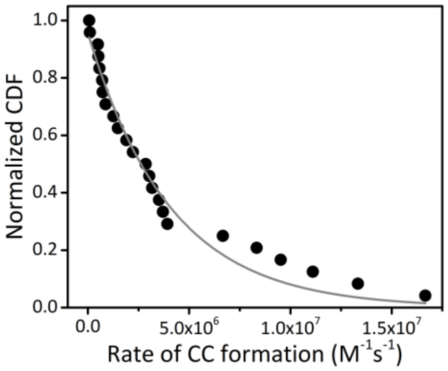


**Fig. S5** Normalized cumulative distribution function (CDF) of initial binding events for each kymograph acquired at low bleaching conditions. Solid line is a fit of mono-exponential decay function.

**Higher force experiments.** As discussed in the main text, we performed experiments at 1 pN and at 5 pN. At 5pN, the RNAP binding profile along the DNA correlates with the local A/T content, similar to the binding profiles that we observed at 1 pN (Fig. S6A). The characteristic event lifetimes (i.e. $t_1$, $t_2$, $t_3$, see Table 1) are directly comparable between the two tensions (fig. S6B). A significant difference is however found in the relative sizes of the populations: the relative number of short events observed at 5 pN (82%) was found to be about twofold larger compared to that at 1 pN (53%) (Fig. S6C). A possible explanation for the increased number of short events (and thus the increased on-rate) is the enhanced signal to noise ratio at higher tension that can result in more effective event detection: increased tension suppresses the thermal fluctuations of the DNA, which reduces motion blurring and thus enhances the signal-to-noise ratio when acquiring fluorescence images at elevated tension. In addition to this effect, the probability to engage into long binding events at promoter sites is reduced at higher tension (cf. Table 1), which may enhance the probability for RNAP to, instead, engage into a shorter-lived non-specific binding mode to the DNA. These results confirm the tension-dependent reduction of promoter recognition that was observed by Harada et al.(4) and are in accordance with the observed tension dependence of promoter binding by the related T7 RNA polymerase by Skinner et al(5). In line with these observations that elevated tension reduces promoter binding, we observe that the position of the longest binding events correlates less well with the promoter locations at high tension than it does at low tension (Fig. S6, C-D).

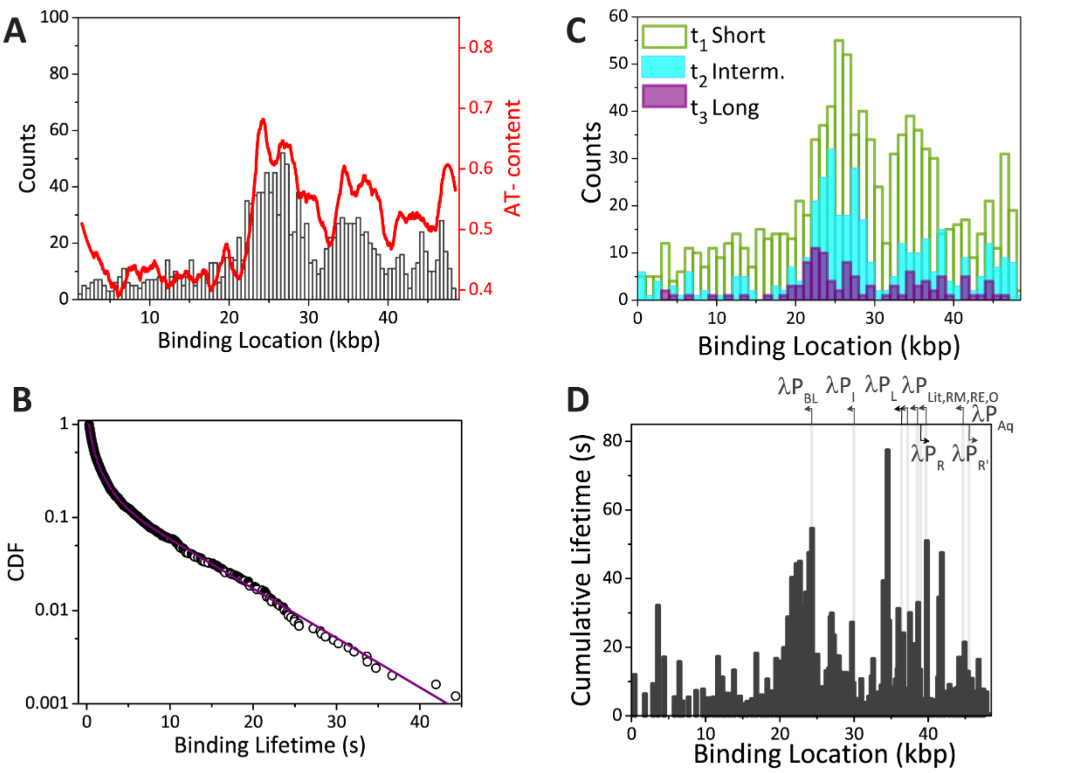


**Fig. S6** High tension experiments (5pN). (A) A histogram of the binding location of RNAP correlates well with the A/T-content along the λ-phage DNA (N=1544). (B) the normalized cumulative distribution function of the binding lifetimes of RNAP to DNA at 5pN can be fit with a three exponential decay that yields off-rates that are directly comparable to those measured at 1 pN tension. (C) Binding location for three populations of binding events that were split according to their event lifetime. (D) Cumulative lifetime as function of binding location along the DNA. Grey lines indicate the locations of promoter sites.

1. Blainey PC, et al. (2009) Nonspecifically bound proteins spin while diffusing along

DNA. *Nat Struct Mol Biol* 16(12):1224–1229.

2. Finn RD, Orlova E V., Gowen B, Buck M, Heel M van (2000) Escherichia coli RNA polymerase core and holoenzyme structures. *EMBO J* 19(24):6833–6844.

3. Friedman LJ, Mumm JP, Gelles J (2013) RNA polymerase approaches its promoter without long-range sliding along DNA. *Proc Natl Acad Sci U S A* 110(24):9740–5.

4. Harada Y, et al. (1999) Single-molecule imaging of RNA polymerase-DNA interactions in real time. *Biophys J* 76(2):709–715.

5. Skinner GM, Kalafut BS, Visscher K (2011) Downstream DNA tension regulates the stability of the T7 RNA polymerase initiation complex. *Biophys J* 100(4):1034–1041.
